# Ground-truth validation of uni- and multivariate lesion inference approaches

**DOI:** 10.1101/2023.03.28.534534

**Authors:** Melissa Zavaglia, Caroline Malherbe, Sebastian Schlaadt, Parashkev Nachev, Claus C Hilgetag

**Author notes:** Correspondence to : Claus C Hilgetag, Institute of Computational Neuroscience, University Medical Center Eppendorf, Hamburg University, Martinistrasse 52, 20246 Hamburg, Germany. These authors contributed equally to the work (Melissa Zavaglia, Caroline Malherbe). Author.

## Abstract

Lesion analysis aims to reveal causal contributions of brain regions to brain functions. Various strategies have been used for such lesion inferences. These approaches can be broadly categorized as univariate or multivariate methods. Here we analysed data from 581 patients with acute ischemic injury, parcellated into 41 Brodmann areas, and systematically investigated the inferences made by two univariate and two multivariate lesion analysis methods via ground-truth simulations, in which we defined a priori contributions of brain areas to assumed brain function. Particularly, we analysed single-region models, with only single areas presumed to contribute functionally, and multiple-region models, with two contributing regions that interacted in a synergistic, redundant or mutually inhibitory mode. The functional contributions could vary in proportion to the lesion damage or in a binary way. The analyses showed a considerably better performance of the tested multivariate than univariate methods in terms of accuracy and mis-inference error. Specifically, the univariate approaches of Lesion Symptom Mapping (LSM) as well as Lesion Symptom Correlation (LSC) mis-inferred substantial contributions from several areas even in the single-region models, and also after accounting for lesion size. By contrast, the multivariate approaches of Multi- Area Pattern Prediction (MAPP), which is based on machine learning, and Multi-perturbation Shapley value Analysis (MSA), based on coalitional game theory, delivered consistently higher accuracy and specificity. Our findings suggest that the tested multivariate approaches produce largely reliable lesion inferences, without requiring lesion size consideration, while the application of the univariate methods may yield substantial mis-localizations that limit the reliability of functional attributions.

## Introduction

Lesion analysis after brain damage is a traditional and powerful approach for revealing causal contributions of brain regions to mental functions. Lesion inferences aim at understanding impaired as well as normal brain function by relating lesion patterns to functional deficits (e.g., cognitive or behavioural impairments) through various strategies, which can be broadly categorized as univariate or multivariate approaches. Univariate approaches assume that the effects of lesions of individual brain region are independent from each other, while multivariate approaches take into account the physical and functional relations among lesion effects of different damaged brain sites.^1,2^ In other words, the univariate approaches provide a statistical comparison of behavioral performance in patients with or without a lesion in an individual region or voxel, whereas the multivariate approaches jointly consider all regions or voxels. Among univariate approaches are Lesion-Symptom Mapping (LSM)^3^ and Lesion Symptom Correlation (LSC).^4^ Most current multivariate methods are based on machine learning classification by support vector machines, such as Multivariate Pattern Analysis (MVPA)^5^ and Multi-Area Pattern Prediction (MAPP).^4^ Similarly, Zhang *et al.*^6^ developed a multivariate lesion symptom mapping approach using a machine learning-based multivariate regression algorithm. An alternative multivariate method is the Multi-perturbation Shapley value Analysis (MSA),^7,8^ which is based on coalitional game theory.

Mah *et al.*^1^ demonstrated that inherent physical dependencies of regions of interest might lead to substantial mis-inferences in univariate analysis methods. While some studies have suggested that univariate methods, specifically LSM, may be able to reduce false positive (mis-)inferences by accounting for total lesion size^9–11^, Inoue *et al*.^12^ found that LSM adjusted for lesion size actually produced a larger bias than LSM without such adjustment. Xu and colleagues^2^ conceptually addressed the problem of the dimensionalities of lesion-deficit mapping, arguing that univariate methods ignore the complexity of the brain in terms of functional interactions among regions, as well as, fundamentally, in terms of anatomical interactions among regions unified by the natural patterns of brain damage. They pointed out that using lesion volume as a regressor in a mass-univariate model would penalise voxels more commonly hit by large lesions and inevitably distort lesion inference.

Given the fundamental role of lesion analysis in neurological research, these findings emphasize the need for a systematic and robust comparison between univariate and multivariate inference approaches (as advocated by Karnath and Smith^13^). In this context, Pustina *et al.*^14^ compared univariate voxel LSM and a multivariate method based on sparse canonical correlation, by using synthetic behaviour as ground truth, and concluded that the multivariate approach produced systematically better accuracy than the univariate method. By contrast, Ivanova *et al.*^15^ compared five univariate and eight multivariate methods (including the method suggested by Pustina *et al.*^14^) using synthetic and real data and saw no clear superiority of a multivariate compared against the univariate methods. Another study by Sperber *et al.*^16^ found that the multivariate method of support vector regression LSM was susceptible to misplacement of statistical topographies along the brain’s vasculature, as is the case for voxel LSM.

Thus, the relative merits of univariate versus multivariate lesion inferences still appear controversial, particularly in the context of large patient cohorts and using complex multivariate methods such as MSA. Therefore, we presently aimed to objectively and systematically compare two univariate (LSM, LSC) and two multivariate (MSA, MAPP) lesion inference methods via ground-truth simulations, analysing data from a large cohort of 581 patients with acute ischemic injury.^1^ Such comparisons cannot be directly performed on clinical datasets, because the true underlying functional contributions and interactions are unknown for such data.^4^ Instead, simulations of ground truth data provide an opportunity to define these contributions and interactions *a priori* and then use a variety of inference approaches to recover the specified mechanisms and compare the accuracy of the respective inferences. The rationale of the study is summarized in Fig. 1.

**Figure 1:**
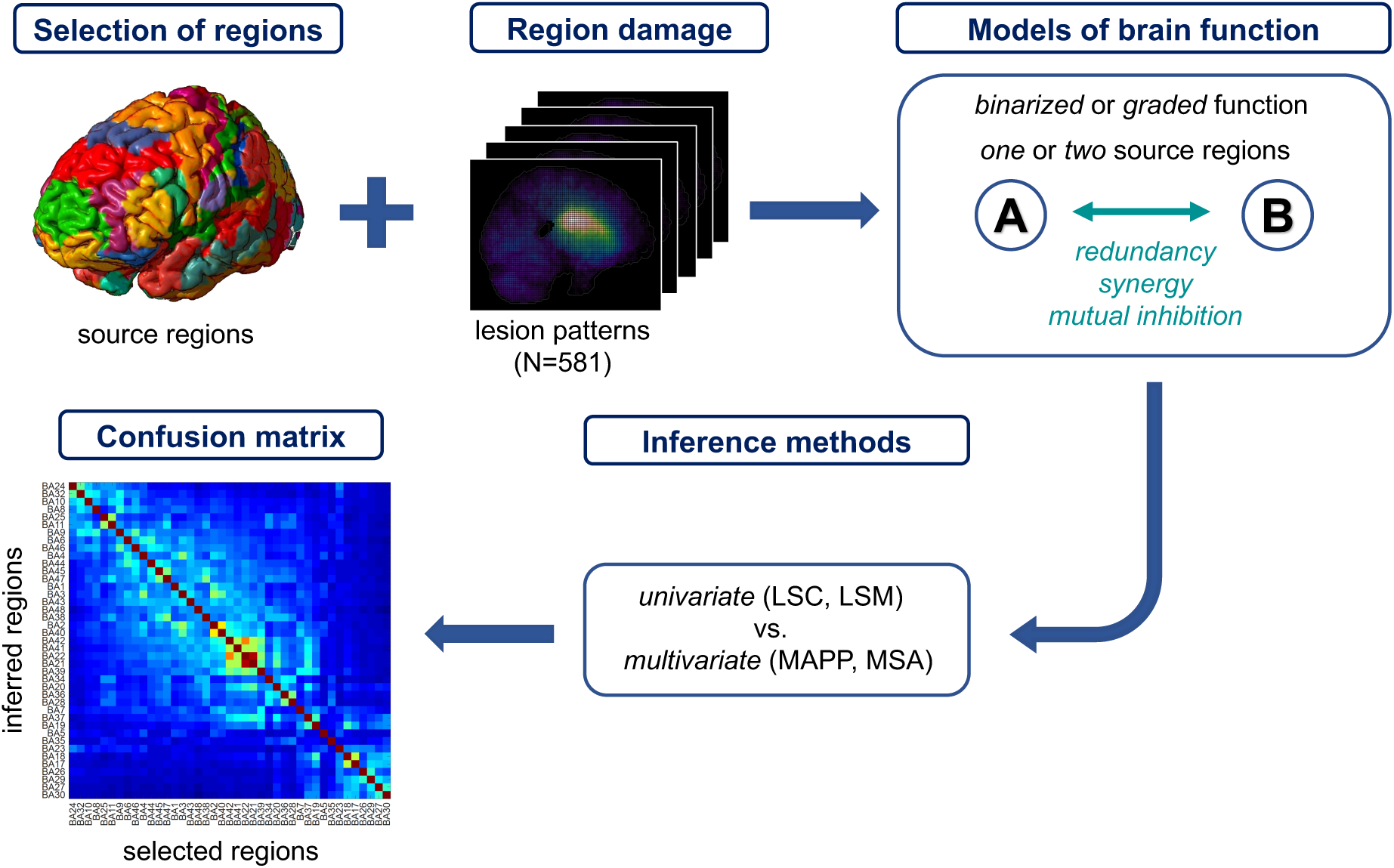
Validation of univariate and multivariate lesion inference methods by ground truth models of brain function. Function was assumed to arise at the regional level, from contributions and interactions of one or two source regions, here selected *a priori* from the set of 41 Brodmann Areas. Region damage was modelled with empirical stroke lesion data (N=581). Damage was assumed to have a categorical or gradual impact on regional function, and regions could interact by synergy, redundancy or mutual inhibition to produce a hypothetical brain function. Lesion patterns and the associated simulated function values were passed to two univariate (Lesion Symptom Correlation, LSC, and Lesion Symptom Mapping, LSM) and two multivariate (Multi- Area Pattern Prediction, MAPP, and Multiperturbation Shapley value Analysis, MSA) lesion inference methods. Finally, the regions inferred by these methods to be functionally contributing were compared with the originally selected regions, determining the accuracy, sensitivity and specificity for each inference method and ground truth model.

Our results confirm the substantially higher accuracy and reliability of the two tested (MAPP and MSA) multivariate compared to the two (LSM and LSC) univariate inferences, also when accounting for lesion size.

## Methods

### Ground-truth simulations at region level

We analysed previously published lesion data from 581 patients with acute ischemic injury,^1^ parcellated into 41 Brodmann areas (BAs), and investigated different lesion models via ground-truth simulations, by defining *a priori* the contributions of brain areas to assumed brain function. For each ground-truth setting, one or two *source regions* were selected as being responsible for a putative brain function, meaning that a theoretical performance score was derived depending on the lesion state of the source regions in each patient’s lesion pattern. In different ground-truth model settings, we explored single versus multiple source region effects and analysed binary versus graded performance datasets.

### Single versus multiple region lesion models

We analysed *single source region* effects, in which only a single area contributed to a presumed brain function as well as *double source region* effects, with two contributing regions. In the single-region setting, a specific lesion model was generated for each of the 41 BA regions selected as a source that was assumed to be solely responsible for a hypothetical brain function. The dataset derived for each of the 41 lesion models was composed of the lesion information for the 581 patients (the same across all models) and the corresponding performance scores, differing from one single-source model to another, depending on which of the 41 BAs was selected as the source of presumed brain function. Differentiating further, the hypothetical performance scores for the 581 patients were derived either in proportion to the intact voxels, or the binary, thresholded state of the specific single or multiple source regions (see subsequent section *Binary versus graded lesion models* for details on the computation of the performance scores). In the double-region models, we focused on pairs of randomly chosen source regions, as well as two specific BAs (BA 39 and BA44) that may be putative loci for visuospatial neglect (cf. Mah *et al*.^1^).

To test if findings on the performance of univariate and multivariate approaches extend to interactions among more than two regions, we also conducted example simulations (described in the Supplemental Material) by considering three regions responsible for a putative score, for the cases of binary and graded redundant triplets. Specifically, the performance was pathological (0) only, if three source regions were lesioned, according to the threshold. An exhaustive analysis of all possible interactions among three regions and their potential functional contributions is beyond the scope of the current study, because of the combinatorial explosion of possible interactions that would have to be considered.^17^

### Binary versus graded lesion models

We explored lesion models that differed in the computation of the performance score in a binary or graded fashion. In the *graded lesion models*, for each ground-truth setting, the functional score was defined in proportion to the level of intactness of the source region. Specifically, intactness for every BA was defined as a relative measure of the intact voxels graded from 0 (complete lesion) to 1 (region completely intact).

In the *binary lesion models*, the binary functional score was defined with respect to a binarization threshold computed as median value of the non-zero lesion size for each region (*median threshold lesion models*), but other thresholds may be used as well (cf. Zavaglia *et al*.^18^ for details on threshold choice). The binary functional score for each ground-truth setting was defined as pathological (0) if the source region was damaged, or normal (1) if the source region was intact, as determined by the threshold.

### Modes of regional interactions

We considered different modes of interactions of the functionally contributing regions in the multiple region models. Specifically, in the case of double region effects and graded lesion models, the performance score was computed in three possible ways: (i) assuming *synergistic* operation, performance was the product of the intactness levels of both source regions; or (ii) assuming *redundant* operation, performance was computed as the average level of intactness of both source regions; or (iii) assuming operation by *mutual inhibition*,^17^ performance was calculated as 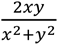, with *x* and *y* the intactness levels of two source regions (performance was set to 1 if the two source regions were completely lesioned (0)).

In the case of double-region effects and binary lesion models, as for the graded lesion models, the performance score was defined in three possible ways: (i) assuming synergistic operation, performance was 1 (normal) only, if both source regions were intact, according to the threshold; (ii) assuming redundant operation, performance was 0 (pathological) only, if both source regions were lesioned, according to the threshold; or (iii) assuming mutual inhibition, performance was 1 (normal) only, if both source regions were lesioned or if both source regions were intact, according to the threshold.

### Incorporation of noise

We also investigated stochastic variants of the *binary lesion model,* for the single source and the double source model with redundancy and with synergy, which added random noise to the simulations. Two principal kinds of noise were considered. The first one occurs if brain regions are intact, yet there is a functional deficit. This kind of noise may represent functional variation across individual patients, potentially related to the impairment of other brain sites. The second kind of noise can occur if regions are lesioned, but the patient does not show a functional deficit. This type of noise may reflect plasticity effects of lesioned regions and compensatory mechanisms across the brain.

Specifically, for the first type of noise, if the single source region was intact according to the threshold, the binary functional score for each ground-truth setting was randomly defined as *1* (normal) only in 90% of the cases. For the second kind of noise, if two sources were lesioned according to the threshold, the binary functional score for each ground-truth setting was randomly defined as 0 (deficit) only in 90% of the cases. In cases of redundancy and synergy we performed analyses where 10% of the functional score was biased and in case of redundancy, we also performed analyses with 50% noise.

### Lesion inference methods

In order to systematically compare univariate and multivariate lesion inference methods, we selected two representative approaches for each category. Specifically, for each generated lesion model (41 single-region lesion models and 50 double-region lesion models), we employed Lesion Symptom Correlation (LSC) and Lesion Symptom Mapping (LSM) as univariate, and Multi-Area Pattern Prediction (MAPP) and Multi-perturbation Shapley value Analysis (MSA) as multivariate analysis methods.

In LSC,^4^ the contributions of the 41 BAs were computed as Spearman rank correlations between all patterns of relative (graded) lesion size and the performance scores (binary or graded, depending on the setting of the models). We used Spearman rank correlations in line with the ordinal nature of the present data.

The second approach, LSM, was originally introduced at voxel level.^3^ As for LSC, we here applied it at the regional level (i.e., for Brodmann Areas). The LSM approach makes use of binary information about the impairment of each BA. The binary dataset was obtained by using a median threshold as explained above. In LSM, we computed the functional contributions of the 41 BAs by statistically comparing via t-test (for the graded simulations) or Mann-Whitney U-test (for the binary simulations) the performance values of all cases in which a particular area was lesioned versus the performance of the cases where the area was intact.

In the MAPP approach,^4^ which is based on machine learning, the contributions of the 41 BAs were obtained by computing the leave-one-out cross-validation via a prediction algorithm with 41 different datasets, obtained respectively by removing each single BA one at a time. The contribution to the prediction error for each BA was obtained as the difference between the Root Mean Square Error (RMSE) computed in the leave-one-out cross validation without the particular BA and the RMSE computed with the complete set of BAs. We used bootstrap^19^ to ensure the robustness of the obtained contributions. Specifically, from the available dataset, we chose 100 random samples with replacement, with the size of the original dataset. We then performed the MAPP approach as outlined above on each of these 100 samples. Finally, the contributions were averaged across the set of 100 samples. Thus, the approach, which is computationally expensive, indicates the ‘importance’ of each BA for the prediction, by quantifying its individual contribution to the prediction error. MAPP makes use of the binary or graded dataset and corresponding binary or graded performance scores depending on the ground-truth settings of the models.

The final approach, MSA,^7^ is a mathematically rigorous method based on game theory for assessing causal function localization from perturbation data. The approach computes an unbiased estimator of the contribution of each brain region from a dataset of multiple lesions. These values represent the regions’ overall contributions to function across all possible brain states. Specifically, the contribution value of a region, formalized as the Shapley value,^20^ represents the importance of the region for the overall behavioral function and it is defined as the difference between the worth of coalitions which contain the region and the worth of coalitions which do not contain it. The MSA variant used here, termed estimated MSA,^21^ is computationally convenient in comparison to the conventional MSA which requires 2*^N^* (*N = number of regions*) performance scores corresponding to the 2*^N^* binary configurations. The estimated MSA can sub-sample orderings and is useful in studies where the number of system elements is too large to enumerate all configurations in a straightforward manner.^21^ We used bootstrap^19^ as described above for MAPP, see Malherbe *et al.*^22^ for using bootstrap with estimated MSA. MSA makes use of a prediction algorithm trained on the available set of configurations, binary or graded, and corresponding performance scores, also binary or graded (depending on the ground-truth setting of the models) to derive performance scores of unknown configurations. For this purpose, in both MAPP and MSA simulations, we used a regression tree predictor (with parameters set to default values). For details on the prediction methodology see Zavaglia *et al*.^18^.

### Data rebalancing

Most of the analyses were faced with imbalanced data, meaning that the different classes were not represented equally in the dataset (e.g., Megahed *et al.*^23^). If a classifier is trained on a dataset in which one of the response classes is rare, it can underestimate the probability of observing a rare event, inducing a bias in the results. To mitigate this imbalance, we trained the binary models for MSA in two ways and used the results with the higher accuracy, performing (i) the bootstrap procedure as usual, or (ii) rebalancing the data by forcing the bootstrap to choose 200 cases in the minority class and the remainder in the majority class. The same procedure was used for the MAPP analysis in each analysis. For the graded models, we used the results with the highest accuracy for the MSA of three possibilities of obtaining the training dataset: (i) the usual bootstrap, (ii) the bootstrap procedure by using 10 classes for the scores instead of the fully graded scores, (iii) the resampling in 10 classes of the score and rebalancing the data during the bootstrap by forcing the bootstrap to choose 200 cases of the minority class and the others in the remaining data. The same procedure was used for the MAPP analysis in each analysis. In both MAPP and MSA simulations, for the single-region binary lesion model with the addition of noise, we used a SVM predictor with linear kernel and cost function = 1. In case of the double source redundancy with noise, we used a regression tree.

### Lesion inference comparison

Ideally, a perfect lesion inference method should infer only one regional contribution in the single region model or two contributions in the double region models, respectively, corresponding to the preselected functional sources. All results from the LSM were corrected for multiple comparisons, according to the Bonferroni correction. For the LSC, we removed all non-significant correlations with the threshold p<0.05. In addition, all results were then normalized. From these corrected normalized results, we calculated the true positive, true negative, false positive and false negative metrics. Specifically, the true positive represents the region(s) expected to be inferred, the false negative the missing expected inferred regions. The true negative regions are the regions that are not expected and not found by the lesion inference method, and the false positive are the found regions that are not expected to be found by the method. As a measure of comparison between the different lesion methods, we computed the accuracy and the weighted accuracy from the sensitivity and specificity and, in addition, the normalized mis-inference error by summing all mis-inferences for a given target (i.e., all inferred regional contributions beyond the ones of the target). For all simulations without noise, we considered all the values for the regions not expected to be found by the method (false positive and true negative), whereas for the simulations with noise, we only considered the regions with a normalized contribution higher than 0.001 to avoid noise-induced spurious results. The threshold of 0.001 was used only for the noisy simulations where we added a stochastic lesion-deficit dependency. For all approaches, this setting induced widespread, very low contributions which were clearly just due to the stochastic setting, but not the analysis approaches themselves. Consequently, the low threshold was used to filter the spurious noise from the actual results in the inferred contributions. All simulations were implemented in MATLAB (The MathWorks Inc.).

### Data availability

The stroke lesion data are available upon request. Scripts for the different inference techniques are available through the original methods papers referenced in the main text. Moreover, there is a recent MSA toolbox developed in our lab available at https://github.com/kuffmode/msa. In addition, scripts for the present analyses are deposited at https://github.com/caromalherbe/MSA-ground-truth

## Results

Fig. 2A presents the location of BAs whose lesions were simulated in single and double target lesion models, and shows the lesion overlap across all 581 cases. Fig. 2B displays the correlation of lesion sizes of the different Brodmann areas across all cases, indicating that many of these areas were affected by the vascular injuries in a related manner.

**Figure 2:**
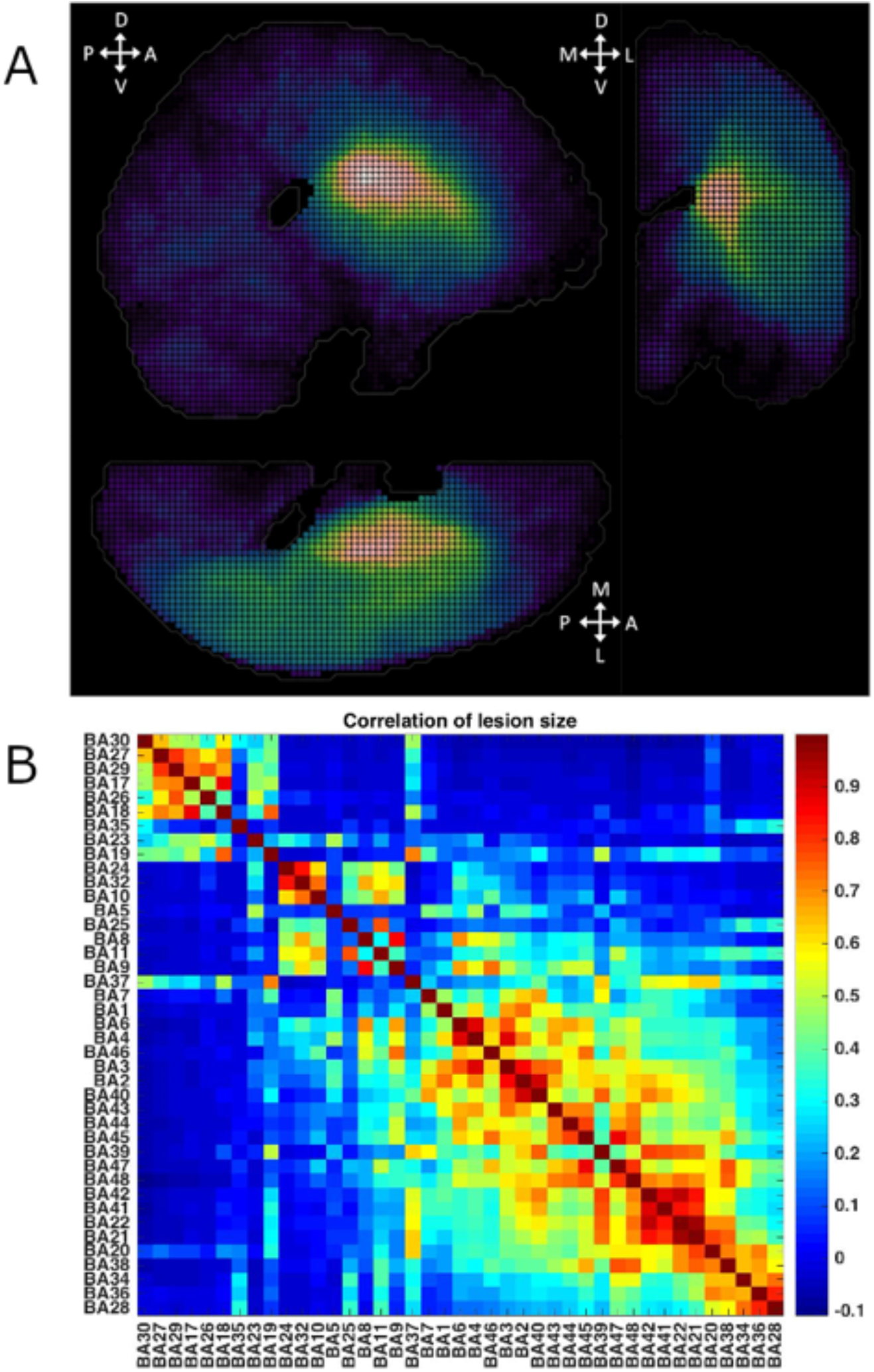
Lesion distribution in the analysed cohort of 581 stroke patients. (A) Overlay plot of stroke lesions from all patients included in the analysis (n=581). The color bar indicates lesion frequency across all patients. (B) Correlation of area lesion patterns. Areas are resorted to show the highest correlation values close to each other. The plot demonstrates that many areas are affected by damage in a similar way.

We performed several evaluations in order to compare lesion inference methods across the various lesion models. Specifically, we compared lesion inferences in terms of accuracy, sensitivity, specificity and mis-inference error, as shown in Table 1 and Table 2. The four inference methods were compared first for the single-region lesion models, using both binary and graded models (results shown in Fig. 3). Then, we investigated the single-region binary lesion model, with the addition of noise (results in Table 1). Finally, we focused on the double-region binary and graded lesion models, both with synergistic, redundant and mutual inhibition performance scores (results for BA39 and BA44 shown in Fig. 4, the remainder listed in Table 1 and Table 2).

**Figure 3:**
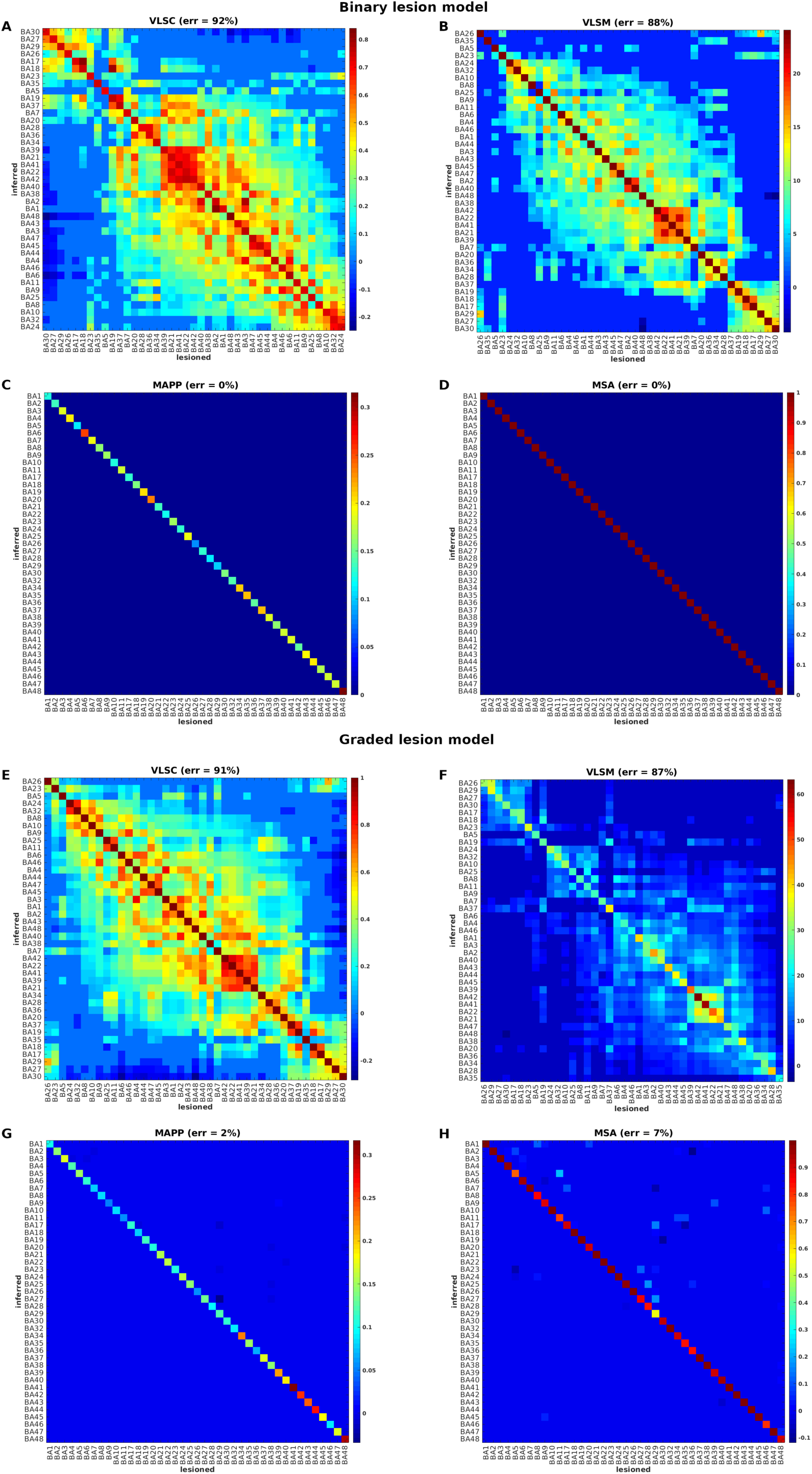
Ground-truth single-region lesion models. Comparison of univariate approaches (panels A, B, E, F) and multivariate approaches (panels C, D, G, H), for ground-truth single-region lesion models. LSC and LSM values were sorted to bring the highest contribution values close to each other. Represented quantities are not normalized.

**Figure 4:**
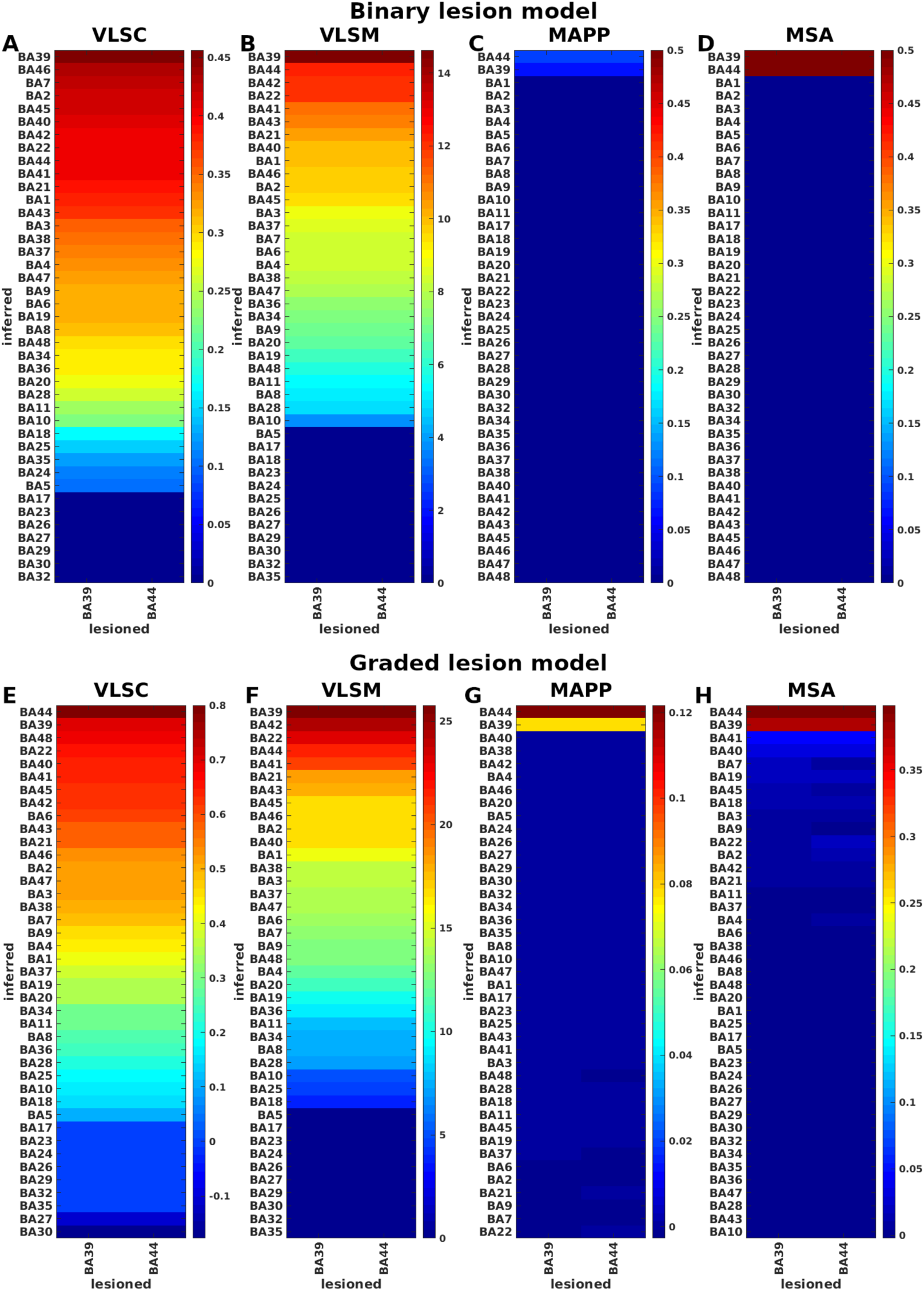
Ground-truth lesion models of double source regions BA39 and BA44. Comparison between univariate (panels A, B, E, F) and multivariate (panels C, D, G, H) methods, for specific ground-truth lesion models of double regions BA39 and BA44. Represented quantities are not normalized.

**Table 1:** Accuracy and misinference errors of uni- and multivariate evaluations of ground-truth models of brain function of single source region and double source regions. LSM_vc: LSM volume corrected; TP: true positive; TN: true negative; FP: false positive; FN: false negative. Accuracy = *(TP+TN)/(TP+TN+FP+FN)*; Accuracy_weighted = *(TP*(1-var(diag))+TN*(1-sum(off_diag)))/(TP+TN+FP+FN)*. Sensitivity = *TP/(TP+FN)*; Specificity = *TN/(TN+FP)*; Mis-inference error = sum of all inferred regional contributions beyond the ones of the target/s. All the quantities are computed on the normalized significant values in the matrices (figures instead show all the values in the matrices, without excluding the nonsignificant values). For the single source binary simulation with noise we used a SVM predictor with linear kernel and cost function = 1, while for all other simulations, we used a regression tree predictor (default parameters). For all simulations without noise, we considered all the values for the regions not expected to be found by the method (FP and TN), whereas for the simulations with noise, we only considered the regions with a contribution higher than 0.001 to avoid all kind of spurious results.

| Inference method | Accuracy % | Accuracy weighted % | Sensitivity % | Specificity % | Misinference error % |
| --- | --- | --- | --- | --- | --- |
| **Single source binary** |  |  |  |  |  |
| LSC (Fig. 2A) | 33 | 5 | 100 | 31 | 92 |
| LSM (Fig. 2B) | 48 | 8 | 100 | 47 | 88 |
| LSM_vc | 71 | 54 | 100 | 70 | 26 |
| MAPP (Fig. 2C) | 100 | 100 | 100 | 100 | 0 |
| MSA (Fig. 2D) | 100 | 100 | 100 | 100 | 0 |
| **Single source graded** |  |  |  |  |  |
| LSC (Fig. 2E) | 29 | 5 | 100 | 27 | 91 |
| LSM (Fig. 2F) | 46 | 8 | 100 | 45 | 87 |
| LSM_vc | 62 | 13 | 100 | 61 | 83 |
| MAPP (Fig. 2G) | 96 | 94 | 100 | 95 | 2 |
| MSA (Fig. 2H) | 95 | 89 | 100 | 95 | 7 |
| **Single source binary and 10% noise** |  |  |  |  |  |
| LSC | 77 | 11 | 100 | 78 | 88 |
| LSM | 74 | 11 | 100 | 73 | 88 |
| MAPP | 98 | 46 | 100 | 98 | 6 |
| MSA | 98 | 73 | 100 | 98 | 4 |
| **Double source binary synergistic** |  |  |  |  |  |
| LSC | 22 | 22 | 100 | 18 | 90 |
| LSM | 35 | 35 | 100 | 32 | 86 |
| MAPP | 100 | 100 | 100 | 100 | 0 |
| MSA | 100 | 100 | 100 | 100 | 0 |
| **Double source binary redundant** |  |  |  |  |  |
| LSC | 39 | 39 | 88 | 37 | 91 |
| LSM | 47 | 47 | 100 | 45 | 86 |
| MAPP | 98 | 98 | 99 | 98 | 1 |
| MSA | 93 | 92 | 100 | 92 | 12 |
| **Double source binary redundant with 10% (or with 50%) noise** |  |  |  |  |  |
| LSC | 80 (79) | 79 (78) | 100 (99) | 79 (78) | 89 (90) |
| LSM | 77 (74) | 76 (74) | 99 (99) | 76 (73) | 86 (86) |
| MAPP | 100 (100) | 100 (100) | 100 (100) | 100 (100) | 95 (96) |
| MSA | 98 (97) | 98 (96) | 100 (97) | 98 (97) | 23 (53) |
| **Double source binary mutual inhibition** |  |  |  |  |  |
| LSC | 41 | 41 | 90 | 40 | 92 |
| LSM | 47 | 46 | 98 | 44 | 85 |
| MAPP | 95 | 95 | 98 | 95 | 1 |
| MSA | 95 | 95 | 96 | 95 | 7 |
| **Double source graded synergistic** |  |  |  |  |  |
| LSC | 19 | 19 | 100 | 15 | 89 |
| LSM | 33 | 33 | 100 | 30 | 87 |
| MAPP | 53 | 53 | 100 | 50 | 8 |
| MSA | 59 | 59 | 100 | 52 | 18 |
| **Double source graded redundant** |  |  |  |  |  |
| LSC | 19 | 19 | 100 | 15 | 89 |
| LSM | 33 | 33 | 100 | 30 | 87 |
| MAPP | 52 | 52 | 100 | 50 | 9 |
| MSA | 64 | 64 | 100 | 62 | 19 |
| **Double source graded mutual inhibition** |  |  |  |  |  |
| LSC | 23 | 23 | 99 | 20 | 89 |
| LSM | 45 | 45 | 89 | 43 | 89 |
| MAPP | 79 | 79 | 68 | 79 | 17 |
| MSA | 94 | 93 | 85 | 94 | 21 |

**Table 2:**
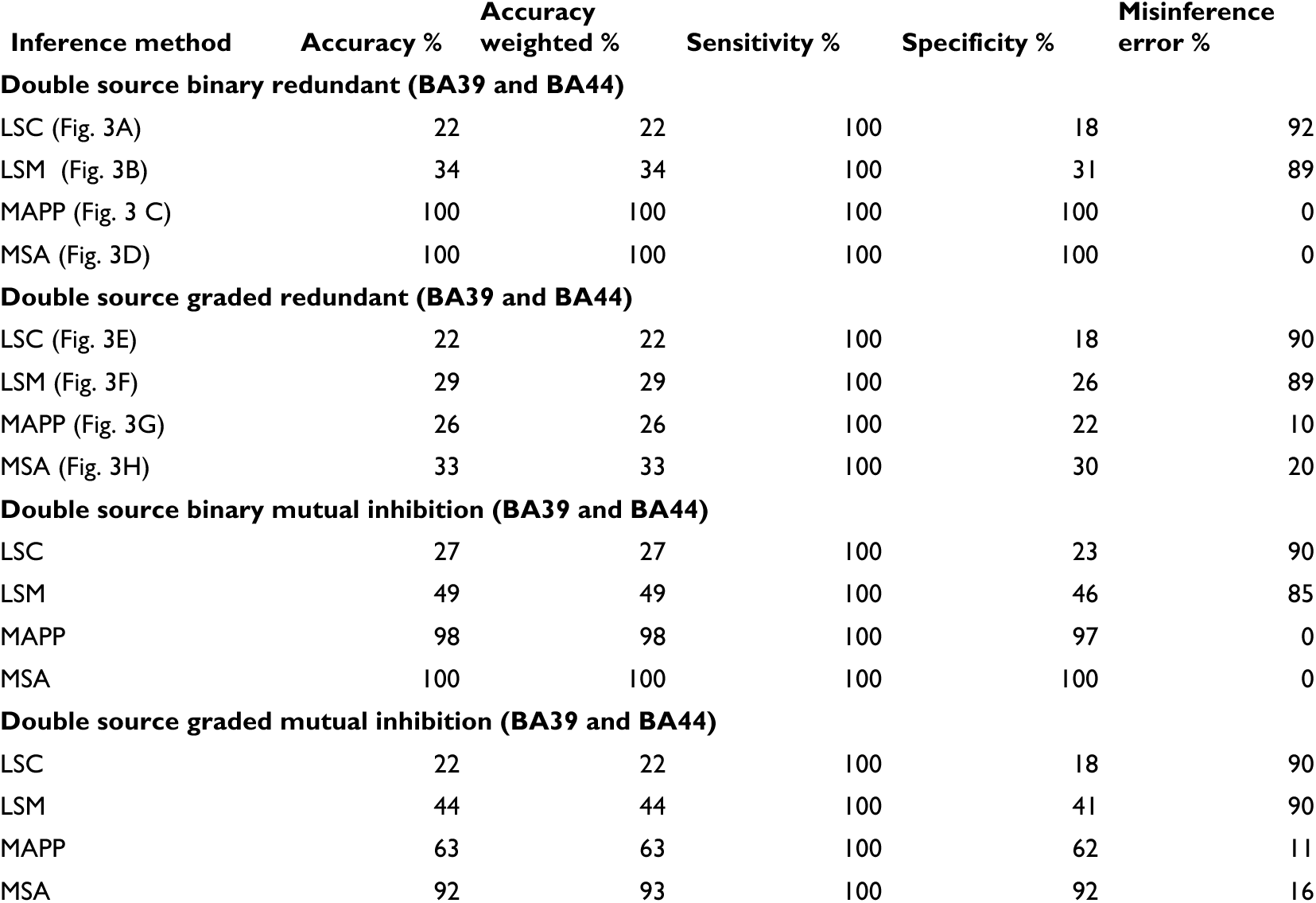
Accuracy and misinference errors of uni- and multivariate evaluations of ground-truth models of brain function for a concrete choice of double source regions: BA39 and BA44. **LSM_vc**: LSM volume corrected; **TP**: true positive; **TN**: true negative; **FP**: false positive; **FN**: false negative. **Accuracy** = *(TP+TN)/(TP+TN+FP+FN)*; **Accuracy_weighted** = *(TP*(1-var(diag))+TN*(1-sum(off_diag)))/(TP+TN+FP+FN)*. **Sensitivity** = *TP/(TP+FN)*; **Specificity** = *TN/(TN+FP)*; **Mis-inference error** = sum of all inferred regional contributions beyond the ones of the target/s. All the quantities are computed on the normalized significant values in the matrices (figures instead show all the values in the matrices, without excluding the nonsignificant values). For all simulations, we used a regression tree predictor (default parameters).

| Inference method | Accuracy % | Accuracy weighted % | Sensitivity % | Specificity % | Misinference error % |
| --- | --- | --- | --- | --- | --- |
| **Double source binary redundant (BA39 and BA44)** |  |  |  |  |  |
| LSC (Fig. 3A) | 22 | 22 | 100 | 18 | 92 |
| LSM (Fig. 3B) | 34 | 34 | 100 | 31 | 89 |
| MAPP (Fig. 3 C) | 100 | 100 | 100 | 100 | 0 |
| MSA (Fig. 3D) | 100 | 100 | 100 | 100 | 0 |
| **Double source graded redundant (BA39 and BA44)** |  |  |  |  |  |
| LSC (Fig. 3E) | 22 | 22 | 100 | 18 | 90 |
| LSM (Fig. 3F) | 29 | 29 | 100 | 26 | 89 |
| MAPP (Fig. 3G) | 26 | 26 | 100 | 22 | 10 |
| MSA (Fig. 3H) | 33 | 33 | 100 | 30 | 20 |
| **Double source binary mutual inhibition (BA39 and BA44)** |  |  |  |  |  |
| LSC | 27 | 27 | 100 | 23 | 90 |
| LSM | 49 | 49 | 100 | 46 | 85 |
| MAPP | 98 | 98 | 100 | 97 | 0 |
| MSA | 100 | 100 | 100 | 100 | 0 |
| **Double source graded mutual inhibition (BA39 and BA44)** |  |  |  |  |  |
| LSC | 22 | 22 | 100 | 18 | 90 |
| LSM | 44 | 44 | 100 | 41 | 90 |
| MAPP | 63 | 63 | 100 | 62 | 11 |
| MSA | 92 | 93 | 100 | 92 | 16 |

Fig. 3 represents the results of the single-region lesion models. Specifically, the panels show the comparison between univariate methods (Fig. 3 A, B, E and F) and multivariate methods (Fig. 3 C, D, G and H), for a single-region median threshold lesion model (binary lesion model) and a single-region graded lesion model. Each column in these confusion matrices represents the 41 contributions of the inferred regions obtained from each ground-truth lesion model (i.e., for each source region which defines a lesion model). Ideally, if a method infers contributions only of the pre-chosen source regions and there are no mis-inferences, the matrices should only have elements on the leading diagonal. The contributions of LSM and LSC were sorted in order to locate the highest values close to the leading diagonal.

The results systematically showed a better performance of multivariate than univariate methods in terms of normalized inference error (reported in percent in Fig. 3, Table 1 and Table 2). Among univariate methods, LSM (accuracy = 48%, specificity = 47%, sensitivity =100% and *err = 80%* for median threshold lesion model and accuracy = 46%, specificity = 45%, sensitivity = 100% and *err = 87%* for graded lesion model) performed better than LSC (accuracy = 33%, specificity = 31%, sensitivity = 100% and *err = 92%* for median threshold lesion model and accuracy = 29%, specificity = 27%, sensitivity = 100% and *err = 91%* for graded lesion model), but both methods mis-inferred substantial contributions from many areas as shown by the clusters outside the leading diagonal. These mis-inferences were attributed to BAs located within the same infarct territory (cf. Fig. 2B). Both multivariate approaches performed perfectly (accuracy = 100%, specificity = 100%, sensitivity = 100% and *err = 0%*) in the median threshold lesion model and almost perfectly in the graded lesion model (MSA: accuracy = 95%, specificity = 95%, sensitivity = 100% and err = 7%; MAPP: accuracy = 96%, specificity = 95%, sensitivity = 100% and err = 2%).

In order to assess the source of the error introduced by univariate approaches, specifically LSM, we quantified the *displacement* from the true location, by computing the total mis-inferences for every ground-truth lesion model across all the inferred BAs, both for graded and median threshold lesion models, and correlated it with (1) the median of *relative* lesion size per area across all cases, (2) the median of *absolute* lesion size per area across all cases and (3) the absolute size per area (details in Supplementary Material). The results suggested that it is the absolute lesion size of regions that matters for the misinference, but not the size of the regions as such. In this context, we also implemented a control for lesion size, as proposed by Sperber and Karnath.^11^ Specifically, we regressed the total absolute lesion size for every case across all BAs on the performance scores. The average correlation between total absolute lesion size and performance scores (across all ground-truth models) was *ρ = -0.34, p = 0.0045* for the median threshold lesion model and *ρ = -0.4,1 p = 0.0044* for the graded lesion model. We performed a linear regression for the graded lesion model and a logistic regression for the median threshold lesion model. We then used the residuals to re-compute the LSM. After adjusting for lesion size (LSM_vc, see Table 1), the accuracy increased to 71 % for the median threshold lesion model and to 62% for the graded lesion model, being larger than in the original LSM analysis, but still substantially smaller than in the multivariate inference approaches.

Fig. 4 represents the same quantities as in Fig. 3, but obtained with the double-region median threshold lesion model and double-region graded lesion model, with redundant performance scores (see also Table 1 and Table 2). The sources chosen for the two regions were BA39 and BA44 (two putative loci for visuospatial neglect^24,25^). The two columns in the matrices represent the 41 contributions of the inferred regions for the two lesion models (i.e., for the two source/target regions). As for the single-region models, among the univariate approaches, LSM (*err = 88%* for median threshold lesion model and *err = 89%* for graded lesion model) performed better than LSC (*err = 92%* for median threshold lesion model and *err = 90%* for graded lesion model), but both methods showed strong mis-inferences. Of the multivariate methods, MSA showed the best performance on the graded lesion model (MSA_*err = 20%* and *MAPP_err = 10%*), generally producing highly reliable lesion inferences, and perfect performance for both MSA and MAPP for median threshold lesion model (err = 0%).

A simulation of the double-region binary redundancy was also performed with different amounts of noise (10% and 50%, respectively) and showed that also under these circumstances the multivariate methods performed better than the univariate ones (Table 1). In the double-region models, in addition to the simulations results reported in Table 1 and Table 2, we also performed simulations by randomly selecting pairs of regions within a group with similar lesion patterns (i.e., with high correlations of lesion size) and within a group with dis-similar lesion patterns (i.e., low correlation of lesion size), to explore the effect of linked lesion patterns of the reliability of lesion inferences. The results in term of accuracy, sensitivity, specificity and mis-inference error are reported in Table S1 (Supplementary Material) and suggest that univariate methods (LSM and LSC) produce mis-inferences independently of whether two areas have similar lesion patterns or not.

In Table S2, we report results for simulations in which three regions were responsible for a putative functional score, interacting redundantly. As in the cases where only one or two regions contributed functionally, we found that the multivariate approaches (MSA and MAPP) performed better than the univariates ones (LSC and LSM). Moreover, simulations with double-region binary synergistic interactions and 10% noise, meaning that two regions were intact but the putative functional score was intact in only 90% of the cases, yielded similar results (Table S2). Moreover, we report in supplemental Table 3S, the effect sizes of our target variables (t-scores, correlation coefficients, prediction error contribution, and normalized contribution values for the approaches of LSM, LSC, MAPP and MSA, respectively) for different setups as well as noise levels. We also compared these values for a lesion inference study of actual clinical data^4^ that used the same inference approaches. The values for the target variables in the present ground truth simulations appear to be in the same range as values seen for the analysis of actual clinical data. ^4^ Moreover, the t-scores in the present analysis (mean values from 3 to 5.5 for all scenarios as reported in the Table 3S) are generally smaller than the ones reported by Pustina and colleagues^14^ in the case of 50% noise (mean value 9.5).

All results reported in Table 1 and Table 2 suggest that the tested multivariate inference methods (MSA and MAPP) consistently perform better (in term of accuracy and mis-inference error) than the univariate methods (LSM and LSC), across all variants of lesion models. MAPP and MSA in most of the cases performed similarly, with a higher computational cost of MAPP than MSA.

## Discussion

Various strategies have been used for inferring brain function from brain lesions in stroke patients. In this study, we systematically compared two univariate (LSM and LSC) and two multivariate (MAPP and MSA) lesion inference approaches via objective ground-truth simulations, by defining *a priori* the contributions of brain areas to assumed brain function. The results systematically showed a better performance of multivariate (MAPP and MSA) than univariate (LSM and LSC) methods in terms of inference error. LSC and LSM inferred substantial contributions for multiple areas even in the single-region models and the mis-inferences were distributed in regions across the infarct territory. Specifically, the strongest mis-inferences were located in BA21, BA22, BA41, BA42 and BA39, corresponding to the lower parietal lobe or temporal lobe, where the largest overlap of lesion cases and correlation of lesion patterns occurs (Fig. 2). The mislocalization across the brain depends on the complex interaction between the multivariate lesion distribution and brain functional architecture^1,2^.

Double-region simulations of two putative loci for visuospatial neglect (BA39 and BA44) showed substantial erroneous displacement of the inferred critical regions with the univariate methods, while no mis-localization was present for the multivariate methods. These results are relevant for correctly attributing functions to neural substrates, such as in the case of inferring the real causal injury location underlying neglect. Mah *et al*.^1^ showed that, with univariate methods, there is a large mis-inference of lesion patterns to the superior temporal gyrus, which might be erroneously considered as critical region for neglect. This region comprises BA22, which is also strongly mis-inferred by the univariate approaches of LSM or LSC in our study.

In the current study, we focused on one or two regions responsible for a putative function, trying to cover systematically many different ways in which they may interact (e.g., redundantly, synergistically or by mutual inhibition, in a graded or binary fashion). We see no principal reason why the present inference methods would not be able to deal with three or more interacting regions responsible for a putative brain function. Along this line, example simulations of three regions contributing functionally and interacting by graded or binary redundancy found similar results as the simulations for single or double source contributions. Specifically, the tested multivariate approaches (MSA and MAPP) performed better than the tested univariate ones (LSC and LSM). Nonetheless, to demonstrate exhaustively that a particular inference approach can deal with three, four or more regions would be much more expensive combinatorially and computationally. For example, three motifs and 13 motifs define all possible relation types between two or three regions, respectively. However, a full characterization of interactions networks composed of four regions already demands 199 motifs, and such numbers would then have to be multiplied by all possible combinations of potential contributions (i.e., three combinations for systems of two regions: just region ‘A’ or region ‘B’ or both ‘A’ and ‘B’ contributing, seven combinations for three regions, etc.), cf. Toba et al. (2020)^17^.

Following recent discussions on the options for decreasing false positives in LSM (c.f. Sperber and Karnath^11^), we also computed LSM contributions after accounting for lesion size, and found a substantial improvement for the median threshold lesion model, but not for the graded lesion model. Generally, however, the mis-inference error with LSM after regressing out total absolute lesion size was still much larger than the errors produced by the tested multivariate approaches (and the accuracy still much smaller than the accuracy for the multivariate approaches). Adding lesion size as covariate is a contested point in the literature. Whereas some authors routinely incorporate lesion size regressions,^11,15^ Xu and colleagues^2^ pointed out the infelicity of adding lesion size as a covariate on principled grounds, as regions of the brain showing strong correlations with lesion volume (i.e., those at the edges of vascular boundaries) will be generally penalised, while others are favoured. More recently, Sperber^26^ and DeMarco and Turkeltaub^27^ argued that a generally valid understanding of the causal relation of lesion size, lesion location, and cognitive deficits is unachievable. They^27^ found a bias for VLSM analyses when the lesion volume was not adequately controlled; however, correcting lesion volume appeared unnecessary when the behavioral variable had no relationship to lesion volume. The authors^27^ conclude that the correction of lesion volume should be specific to each research question. Our empirical analyses particularly underline the fact that lesion size regression is not an automatic cure for the imperfections of the tested univariate lesion inference (LSM).

While perfect noiseless lesion-deficit associations are certainly not realistic, it should be pointed out that even without noise, the tested univariate methods did not provide a perfect performance in predicting a lesioned target region. Thus, the comparatively high performance of the tested multivariate methods was not just due to the absence of noise. Naturally, lesion inference approaches should generally strive to apply the best available analysis technique to data that contains as little noise as possible (cf. Toba et al.^28^). Nonetheless, our simulations with noise in single-region and double-region settings also showed a higher performance of the tested multivariate approaches compared to the univariate ones.

Finally, we also compared the size of the respective target variables in the four inference approaches across the different model scenarios as well as with the size of the same variables in an analysis of actual clinical data.^4^ The comparison (Table S3) showed that the effect sizes varied only moderately across the scenarios without noise or with 10% or 50% added noise. Generally, t-scores of the simulations in the present study were smaller than t-scores in the ground truth simulations by Pustina et al. when they used 50% noise or less. Moreover, the size of the present target variables was similar to those for actual clinical data that were analyzed with the same inference approaches^4^. Naturally, a direct comparison between the analysis of clinical data and ground truth data is difficult, because the actual ground truth for the clinical data is unknown and the assumption that clinical data simply consist of the kind of ground truth considered here with a superposition of some noise may not hold.

Generally, our findings confirm the presence of substantial mis-inferences of locations in the tested univariate lesion analysis approaches (LSM and LSC), and demonstrate that the tested multivariate approaches consistently produced highly reliable lesion inferences, without requiring lesion size correction. In particular, the game-theoretical MSA generally provided a good performance, combining high accuracy and low computational cost. Nonetheless, future work ought to expand these findings by testing even more intricate models of brain function^17^ and lesion effects, in order to derive a comprehensive understanding of lesion inference approaches.

## Supplementary material

Supplementary material is available at *Brain Communications* online.

